# Trimeric SARS-CoV-2 Spike interacts with dimeric ACE2 with limited intra-Spike avidity

**DOI:** 10.1101/2020.05.21.109157

**Authors:** Irene Lui, Xin X. Zhou, Shion A. Lim, Susanna K. Elledge, Paige Solomon, Nicholas J. Rettko, Beth Shoshana Zha, Lisa L. Kirkemo, Josef A. Gramespacher, Jia Liu, Frauke Muecksch, Julio Cesar Cetrulo Lorenzi, Fabian Schmidt, Yiska Weisblum, Davide F. Robbiani, Michel C. Nussenzweig, Theodora Hatziioannou, Paul D. Bieniasz, Oren S. Rosenburg, Kevin K. Leung, James A. Wells

## Abstract

A serious public health crisis is currently unfolding due to the SARS-CoV-2 pandemic. SARS-CoV-2 viral entry depends on an interaction between the receptor binding domain of the trimeric viral Spike protein (Spike-RBD) and the dimeric human angiotensin converting enzyme 2 (ACE2) receptor. While it is clear that strategies to block the Spike/ACE2 interaction are promising as anti-SARS-CoV-2 therapeutics, our current understanding is insufficient for the rational design of maximally effective therapeutic molecules. Here, we investigated the mechanism of Spike/ACE2 interaction by characterizing the binding affinity and kinetics of different multimeric forms of recombinant ACE2 and Spike-RBD domain. We also engineered ACE2 into a split Nanoluciferase-based reporter system to probe the conformational landscape of Spike-RBDs in the context of the Spike trimer. Interestingly, a dimeric form of ACE2, but not monomeric ACE2, binds with high affinity to Spike and blocks viral entry in pseudotyped virus and live SARS-CoV-2 virus neutralization assays. We show that dimeric ACE2 interacts with an RBD on Spike with limited intra-Spike avidity, which nonetheless contributes to the affinity of this interaction. Additionally, we demonstrate that a proportion of Spike can simultaneously interact with multiple ACE2 dimers, indicating that more than one RBD domain in a Spike trimer can adopt an ACE2-accessible “up” conformation. Our findings have significant implications on the design strategies of therapeutic molecules that block the Spike/ACE2 interaction. The constructs we describe are freely available to the research community as molecular tools to further our understanding of SARS-CoV-2 biology.

## Introduction

In late 2019, a novel, pathogenic coronavirus (SARS-CoV-2) entered the human population and has since spread throughout the world. The number of people suffering from the associated disease (COVID-19) continues to rise, increasing the need for effective therapeutic interventions. SARS-CoV-1 and SARS-CoV-2 Spike proteins are highly homologous (~76% sequence identity). Similar to SARS-CoV-1, the interaction between the SARS-CoV-2 Spike protein and the angiotensin-converting enzyme 2 (ACE2) on human cells is critical for viral entry into host cells (Gralinski & Menachery, 2020; Tai et al., 2020; Wu et al., 2020). SARS-CoV-2 Spike is an obligate trimer, while ACE2 presents as a dimer on the cell surface (Chen, Liu, & Guo, 2020). Several high-resolution structures of SARS-CoV-2 Spike receptor binding domain (Spike-RBD) bound to ACE2 have been published (Lan et al., 2020; Yan et al., 2020). However, as of this writing, structures of SARS-CoV-2 Spike trimer in complex with either the dimeric or monomeric form of ACE2 have not been reported, resulting in an incomplete understanding of the nature of this interaction.

Structural studies of trimeric SARS-CoV-2 and SARS-CoV-1 Spike protein demonstrate that each of the Spike-RBDs, as in other coronaviruses, can undergo hinge-like movements to transition between “up” or “down” conformations. The host ACE2 receptor can only interact with an RBD in the “up” conformation, whereas the “down” conformation is inaccessible to ACE2 (Wrapp, Vlieger, et al., 2020). The RBDs of SARS-CoV-1 Spike can rotate away from the “down” position by different angles to an “up” position (Gui et al., 2017). Several cryo-EM structures report that approximately half of the SARS-CoV-1 and 2 Spike particles are in the “three-down” closed conformation and half in the “one-up” open conformation (Song, Gui, Wang, & Xiang, 2018; Walls et al., 2020), while another cryo-EM study on SARS-CoV-1 Spike reported 39% of Spike in the “two-up” conformation and 3% in the “three-up” conformation (Kirchdoerfer et al., 2018). The different conformations of Spike observed by these cryo-EM studies may be affected and/or limited by the properties of the grid and sample preparation conditions, and may also reflect differences between SARS-CoV-1 and 2. Thus, it remains unknown how many RBDs of SARS-CoV-2 are accessible within a trimeric Spike to bind ACE2 under physiological conditions, and thus the degree to which intra-Spike avidity plays in the interaction of ACE2 with SARS-CoV-2 Spike. Recombinant ACE2 and an engineered dimeric ACE2-Fc fusion have been shown in several studies to neutralize SARS-CoV-2 virus (Lei et al., 2020; Li et al., 2020; Monteil et al., 2020). However, it remains unknown whether the dimeric form of ACE2 offers any affinity enhancements through avidity compared to a monomer. Understanding the role and mechanism of intra-Spike avidity in binding is important for engineering tight binding antagonists to neutralize virus infection.

To elucidate the nature of the interaction between dimeric ACE2 and trimeric SARS-CoV-2 Spike, we performed a thorough characterization of the binding interactions of the different multimeric forms of Spike-RBD and ACE2 **(Figure 1A)**. The results reveal that while both the ACE2 monomer and ACE2-Fc dimer can bind the isolated Spike-RBD, only the ACE2-Fc dimer can bind tightly to the trimeric Spike ectodomain (FL-Spike). Interestingly, the affinity of the ACE2-Fc dimer is much higher to the RBD-Fc dimer than to FL-Spike, suggesting that although intra-molecular avidity plays a role in both interactions, its effect is compromised in the context of FL-Spike. Consistent with this, we find that ACE2 associates more slowly to FL-Spike, which indicates that the RBDs in FL-Spike protein are not readily accessible to ACE2. To further probe the conformational landscape of the RBDs in FL-Spike, we engineered ACE2-Fc as split-Nanoluciferase (NanoLuc) reporters. We found that a proportion of FL-Spike can interact with multiple ACE2-Fc molecules simultaneously, indicating that more than one RBD domain in a Spike trimer can adopt an “up” conformation. Using pseudotyped and SARS-CoV-2 virus neutralization assays, we further show that ACE2-Fc dimer is much more potent at neutralizing virus than ACE2 monomer. These findings extend our biochemical insight into how a Spike trimer binds to an ACE2 dimer, and have important implications for how multimeric therapeutic molecules, such as dimeric ACE2 or antibody-based biologics, can effectively target SARS-CoV-2.

**Figure 1.**
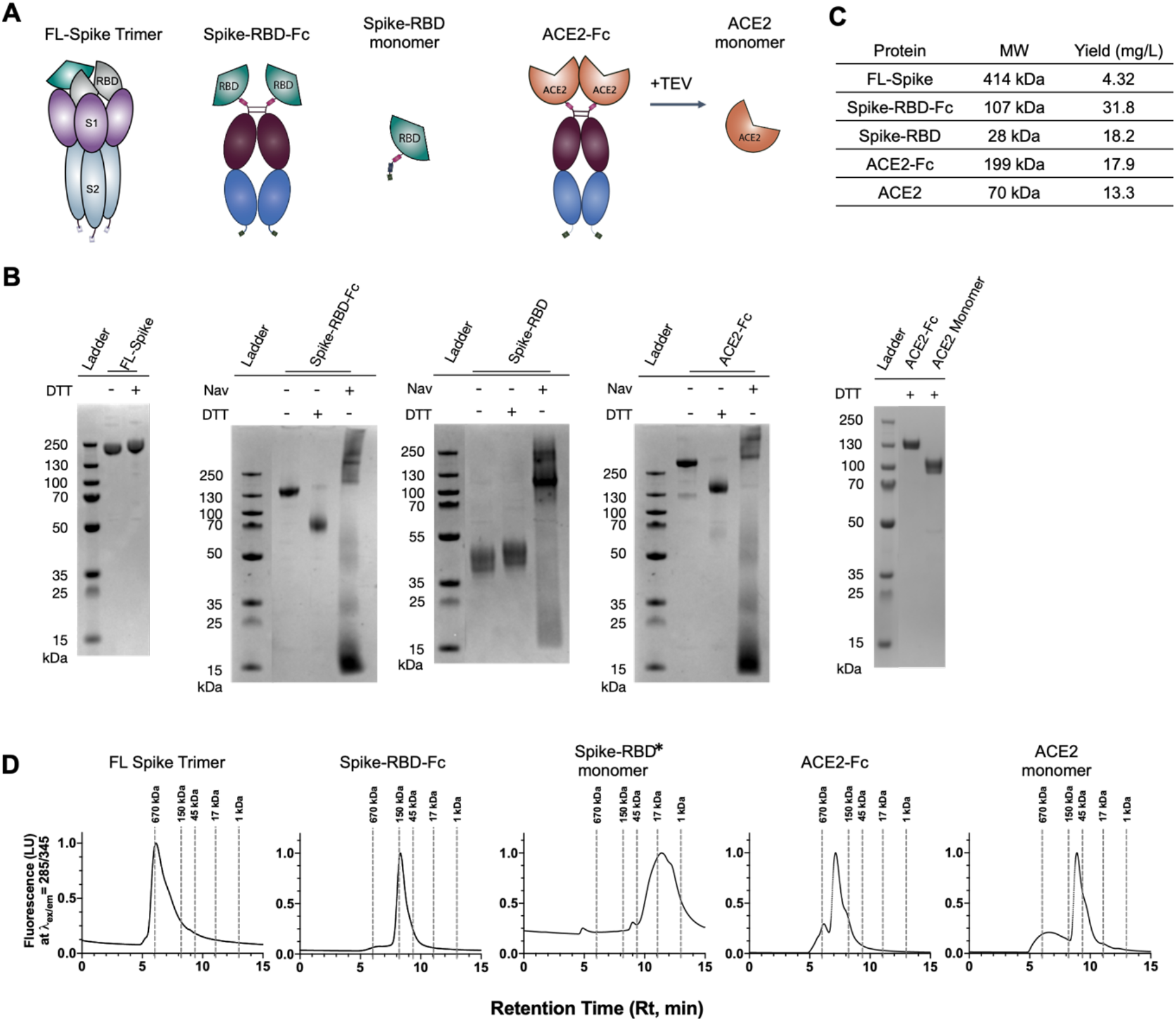
Purification and characterization of Spike and ACE2 variants. **(A)** Cartoon representation of antigens: FL-Spike, Spike-RBD-Fc, Spike-RBD, ACE2-Fc and ACE2. **(B)** SDS-PAGE gel (4-12%) analysis of purified proteins stained with Coomassie blue. Protein was incubated with reducing agent or neutravidin to determine biotinylation. **(C)** Purification method and expression yield per construct. **(D)** SEC traces of purified protein. Dotted lines mark retention time of a molecular weight standard. *Retention time of Spike-RBD monomer does not precisely match elution profile of standard, but further SDS-Page gel analysis of SEC fractions shows pure protein at the correct molecular weight **(Supplementary Figure 1)**.

## Results

### Design and generation of different multimeric Spike/ACE2 proteins

To study the interaction between ACE2 and SARS-CoV-2 Spike, we constructed a panel of Spike and ACE2 proteins in various multimeric formats **(Figure 1A)**. Trimeric SARS-CoV-2 Spike ectodomain (aa 1-1213) (FL-Spike) was expressed and purified as described (Amanat et al., 2020). The construct generously provided by the F. Krammer Lab has the furin cleavage site removed, a pair of stabilizing mutations and a C-terminal T4 trimerization motif followed by a 6xHis tag added. These modifications have been widely used for structural analysis (Gui et al., 2017; Kirchdoerfer et al., 2018; Song et al., 2018; Walls et al., 2020; Wrapp, Wang, et al., 2020). We designed a dimeric form of the Spike Receptor Binding Domain (aa 328-533) (Spike-RBD-Fc) containing a TEV-cleavable Fc-fusion molecule with a C-terminal Avi tag for biotinylation (Czajkowsky, Hu, Shao, & Pleass, 2012). We also generated a monomeric form of Spike-RBD (aa 328-533) (Spike-RBD-monomer) with a C-terminal TEV-8xHis-Avi. The human ACE2 ectodomain contains a N-terminal peptidase domain (aa 18-614) and a C-terminal dimerization domain (aa 615-740). We designed the monomeric form of ACE2 (aa 18-614) (ACE2-monomer) with a C-terminal TEV-8xHis-Avi tag, and the dimeric form of ACE2 (aa 18-614) (ACE2-Fc) as a TEV-cleavable Fc-fusion molecule with a C-terminal Avi tag (Czajkowsky et al., 2012). All of the ACE2 and Spike proteins were expressed in BirA-ER-expressing Expi293 cells (Howarth et al., 2008; Martinko et al., 2018).

The Fc-fusion molecules were purified by Protein A affinity chromatography, and the Spike-RBD-monomer and FL-Spike by Ni-NTA affinity chromatography (**Figure 1B, 1C**). However, the ACE2 monomer did not express but was generated instead by TEV release from ACE2-Fc (**Figure 1A, 1B**). All these proteins, except the ACE2 monomer were >95% biotinylated during expression (**Figure 1B**), facilitating their use on avidin-functionalized surfaces and beads. Size exclusion chromatography was performed to confirm the oligomerization state of the different proteins. FL-Spike, Spike-RBD-Fc, ACE2 monomer, and ACE2-Fc all eluted at their expected elution times (**Figure 1D**), indicating successful generation of the different multimeric forms of Spike and ACE2 proteins. The Spike-RBD-monomer eluted later than expected, but further analysis of the associated SEC fractions by SDS-PAGE showed the pure protein at the correct molecular weight (**Supplementary Figure 1**). Differential scanning fluorimetry (DSF) of ACE2-Fc and ACE2 monomer showed these two proteins had similar Tm values **(Supplementary Figure 2)**.

### ACE2 dimerization is important for binding to the trimeric SARS-CoV-2 Spike

To understand how oligomerization affects Spike/ACE2 interaction, we determined the affinity and binding kinetics of the different Spike-RBD and ACE2 proteins by bio-layer interferometry (BLI) (**Figure 2**). Spike-RBD-monomer, Spike-RBD-Fc and FL-Spike were immobilized on Streptavidin or Ni-NTA sensors, and allowed to bind ACE2 monomer or ACE2-Fc in solution. A four-fold avidity effect was observed for the Spike-RBD-monomer/ACE2-Fc interaction (K_D_ = 5.5 nM, **Figure 2D**) compared to the Spike-RBD-monomer/ACE2-monomer interaction (K_D_ = 22.4 nM, **Figure 2A**). In contrast, the Spike-RBD-Fc/ACE2-Fc interaction (K_D_ < 10^−12^ M, **Figure 3E**) showed >1000-fold increase in binding affinity compared to the Spike-RBD-Fc/ACE2-monomer interaction (K_D_ =13.2 nM, **Figure 2B**). This dramatic increase in affinity is driven by a massive decrease in the off rate and little change in on rate, which indicates a strong intramolecular two-on-two avidity between Spike-RBD-Fc and ACE2-Fc.

**Figure 2.**
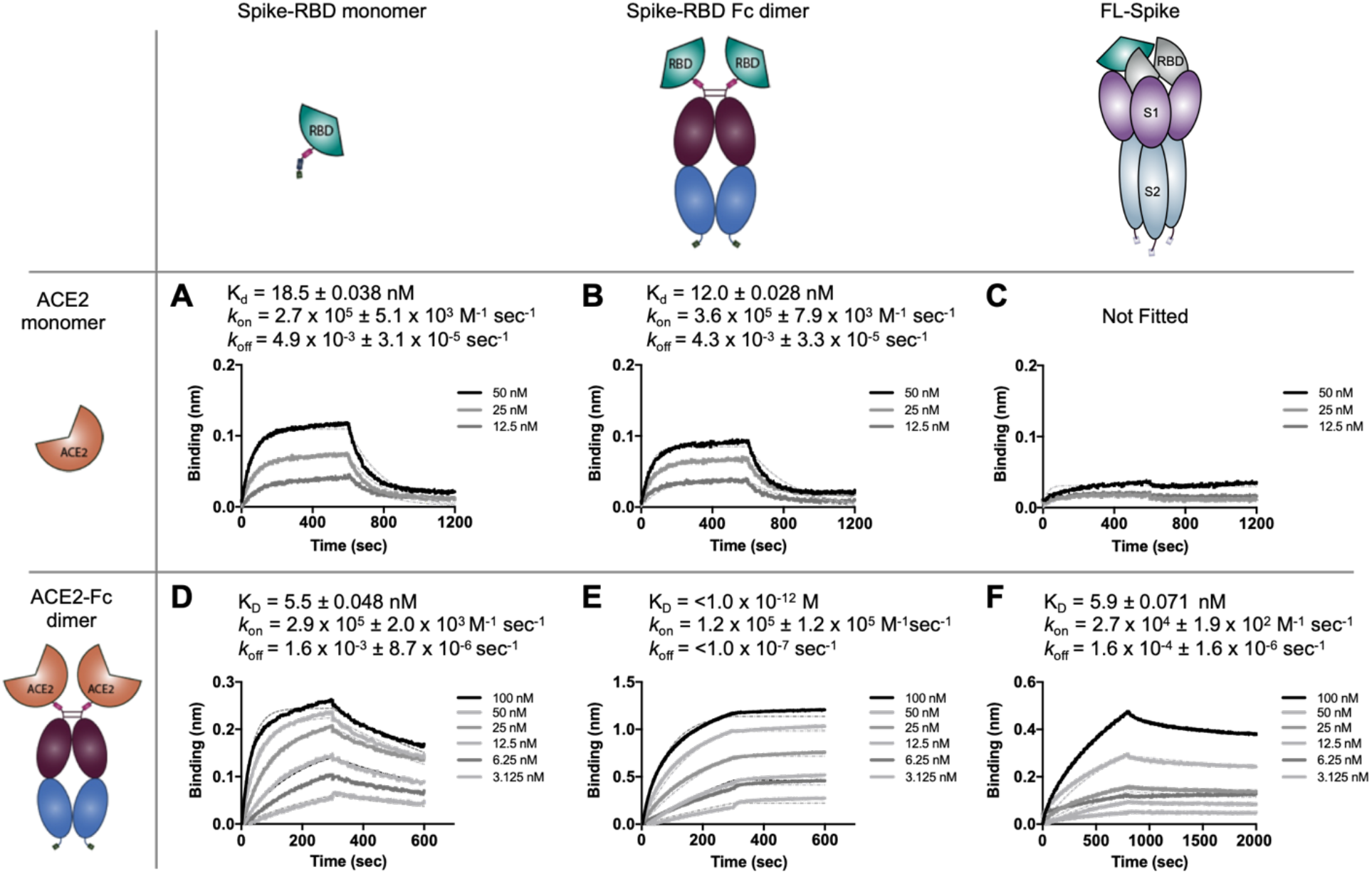
BLI characterization of binding affinity and kinetics of Spike and ACE variants. Bio-layer Interferometry (BLI) data of ACE2 monomer or ACE2-Fc dimer binding to immobilized Spike RBD-monomer, RBD-Fc dimer, or trimeric FL-Spike ectodomain. Spike variants were loaded onto sensors until binding was between 0.4-0.6 nm. ACE2 monomer or ACE2-Fc were used as analytes in solution, with concentrations ranging from 100 nM to 3.125 nM. Although ACE2 monomer bound Spike-RBD-monomer and Spike-RBD-Fc at K_D_ of 18.5 and 12.0 nM, respectively, it did not bind FL-Spike strongly **(A-C)**. In contrast, the ACE2-Fc dimer bound to these forms with affinities of 5.5 nM, <1 pM, and 5.9 nM **(D-F)**. Although ACE2-Fc can bind Spike-RBD-Fc with strong intramolecular avidity, this 2-on-2 interaction is not present in the context of FL-Spike, indicating that only one arm of ACE2-Fc can engage an RBD in FL-Spike. The decreased *k*_on_ and *k*_off_ of ACE2-Fc in the context of FL-Spike **(F)** compared to Spike-RBD-monomer **(D)** indicates that conformational changes of the RBD in the context of the Spike trimer are important in the interaction of ACE2-Fc to FL-Spike.

**Figure 3.**
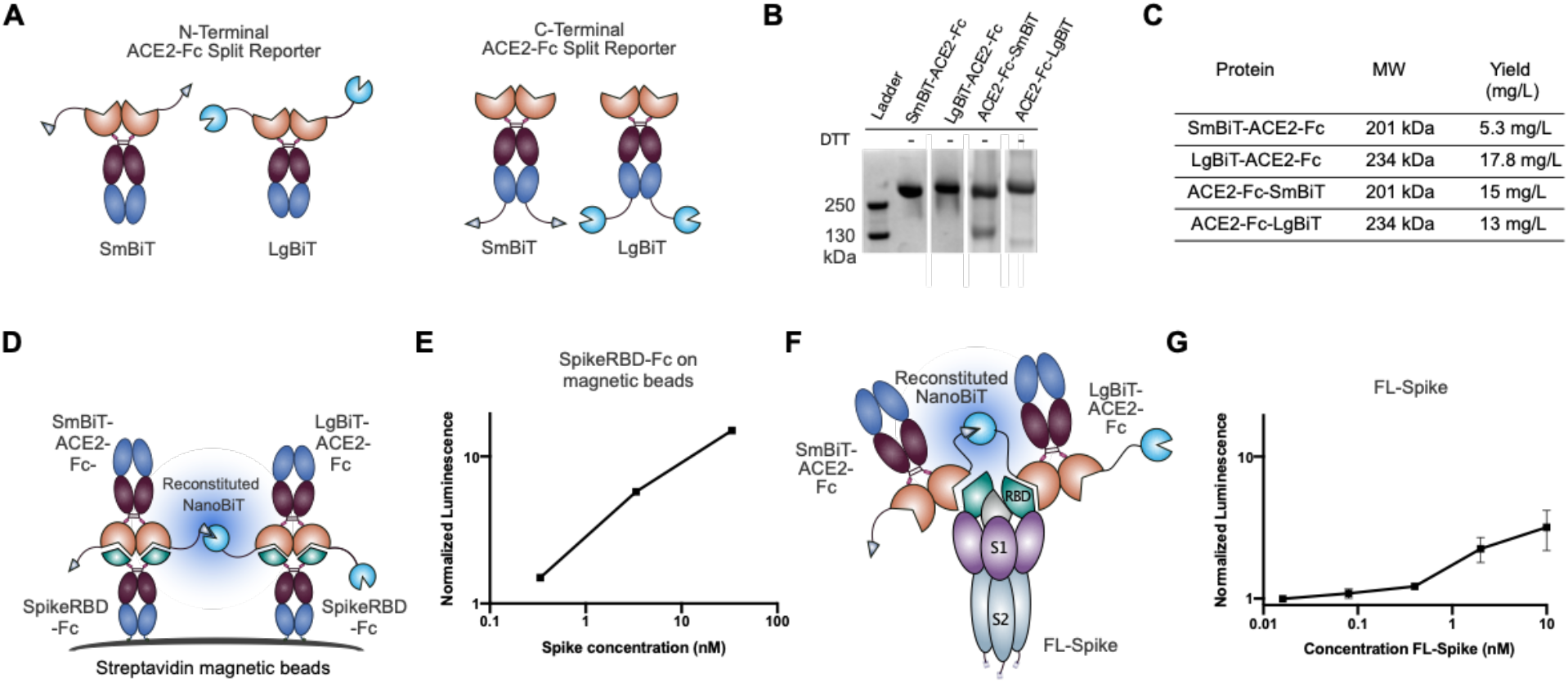
ACE2-Fc split luciferase experiments demonstrates more than one RBD in FLSpike are available to bind ACE2. **(A)** Cartoon depiction of the N-terminal and C-terminal ACE2 NanoBiT split luciferase constructs. **(B)** SDS-PAGE gel showing expression and purification of ACE2 fusions. All show a major band at the expected molecular weight of the dimer. **(C)** All of the ACE2 split-reporter fusions had good yields (>5 mg/L). **(D)** Cartoon depiction of the N-terminal ACE2 NanoBiT sensor system to detect Spike-RBD-Fc bound to streptavidin magnetic beads. **(E)** The N-terminal ACE2 NanoBiT system is able to detect Spike-RBD-Fc bound magnetic beads with better sensitivity than the C-terminal fusions **(Supplementary Figure S4)**. **(F)** Cartoon depiction of the N-terminal ACE2 NanoBiT sensor system to detect FL-Spike in solution. **(G)** The N-terminal ACE2 NanoBiT system is able to detect FL-Spike in solution, indicating that more than one RBD in a FL-Spike trimer can be in the “up” conformation for two ACE2-Fc molecules to bind and reconstitute the split enzyme. For all assays, the luminescence signal is normalized to a no-bead control and the average and standard deviation is plotted (N=3).

We next tested binding of ACE2 monomer and ACE2-Fc to FL-Spike to determine the affinity and avidity effect. Surprisingly, we found that the binding interaction between ACE2 monomer and FL-Spike is very weak and could not be measured accurately (**Figure 2C**). By contrast, the ACE2-Fc interacts with FL-Spike with a K_D_ of 5.9 nM, similar to the affinity for the isolated Spike-RBD-monomer (**Figure 2F**). This suggests that the presence of two ACE2 molecules in close proximity in ACE2-Fc is essential for a productive interaction with FL-Spike, and that a single ACE2 monomer is not sufficient to bind an RBD on FL-Spike. However, the affinity between FL-Spike/ACE2-Fc interaction was substantially less than Spike-RBD-Fc/ACE2-Fc interaction (K_D_ < 10^−12^ M, **Figure 2E**), suggesting that the high-avidity two-on-two interaction is compromised in the context of FL-Spike. This could be due to geometric/steric constraints or the unique conformations of the RBDs in the FL-Spike context. When FL-Spike is loaded to a much higher density on the BLI sensor (load to 2.0 nm) and probed with ACE2-Fc, we see that avidity can be recovered **(Supplementary Figure S3A)**. This indicates that if Spike is presented at high density, the ACE2-Fc arms can engage two RBDs if neighboring Spike trimers are close enough. In contrast, monomeric ACE2 did not bind FL-Spike strongly even when FL-Spike was loaded until saturation, further demonstrating the importance of ACE2 dimerization for interacting with FL-Spike **(Supplementary Figure S3B)**.

Interestingly, the *k*_on_ of the FL-Spike/ACE2-Fc interaction (**Figure 2C and 2F**) is ~10-fold lower than the interactions between Spike-RBD-monomer or Spike-RBD-Fc with ACE2 or ACE2-Fc (**Figure 2A, 2B, 2D, and 2E**), while the *k*_off_ is also ~10-to 20-fold lower. The decreased *k*_on_ suggests that the RBDs in FL-Spike protein may have to undergo a conformational change for binding to ACE2. Previous cryo-EM studies on SARS-CoV-2 FL-Spike show that approximately half of the particles have the three RBD domains in the “down” conformation (Walls et al., 2020). Molecular dynamics simulations of SARS-CoV-2 FL-Spike suggest that the RBD exists in a series of conformations between the “down” and “up” states, and any RBD with an angle lower than 52.2° from the body of the trimer are inaccessible to ACE2 (Peng et al., 2020). Our data supports that a significant proportion of the RBDs in FL-Spike protein are in a “closed” or partially “closed” state inaccessible to ACE2, and the RBD has to open up to allow binding to ACE2. The decreased *k*_off_, on the other hand, suggests that the presence of multiple RBDs within the context of a FL-Spike could slow down the dissociation of ACE2-Fc.

### Split reporter assays indicate more than one RBD in a Spike trimer can be in the “up” conformation when it binds ACE2

To further investigate the RBD conformational landscape in FL-Spike, we designed a split-NanoLuc system to orthogonally probe the Spike/ACE2 interaction. Split-NanoLuc enzymes, in particular NanoBiT (Promega), have been broadly used to detect protein-protein interactions and to study analyte concentrations (Dixon et al., 2016). The NanoBiT system is composed of LgBiT and SmBiT. SmBiT is an 11 amino acid peptide which has a low intrinsic affinity to LgBiT (K_D_ = 190 μM), but when SmBiT and LgBiT are in close proximity, the two subunits assemble to form an active luciferase enzyme. To interrogate the interaction between ACE2 and Spike, we engineered ACE2-Fc reporter molecules where SmBiT or LgBiT were fused at the N- or C-termini (**Figure 3A**). All constructs expressed at high yield and purity **(Figure 3B, 3C)**.

To functionally validate the split reporter system, we immobilized Spike-RBD-Fc on streptavidin magnetic beads at high-density. Incubation with 1 nM of ACE2-Fc-SmBiT and ACE2-Fc-LgBiT, or 1 nM of SmBiT-ACE2-Fc and LgBiT-ACE2-Fc with substrate showed dose-dependent luminescence signal, consistent with an assembled functional split enzyme reporter and intermolecular proximity (**Figure 3D, 3E**). We found the N-terminal fusion reporter pair showed higher sensitivity compared to the C-terminal fusion reporter pair (**Supplementary Figure S4**) suggesting that the increased entropy from the flexible linker and Fc domain reduces productive luciferase reconstitution.

We next used these split reporters to interrogate the soluble FL-Spike trimer (**Figure 3F, 3G**). Increasing concentrations of soluble FL-Spike were incubated with ACE2-Fc-SmBiT/LgBiT or SmBiT/LgBiT-ACE2-Fc, followed by the addition of substrate. The SmBiT/LgBiT-ACE2-Fc reporters showed dose-dependent increase in luminescence signal with 1-10 nM FL-Spike. This result suggests that although FL-Spike cannot form a high-avidity two-on-two interaction with both arms of ACE2-Fc at a time, there is a proportion of FL-Spike proteins with “two-up” or perhaps even “three-up” RBDs that can simultaneously interact with multiple ACE2-Fc molecules. We cannot resolve if these “two-up” or “three-up” conformations are present prior to ACE2 binding, or appear because ACE2 binding induces a conformational change, which enables two or more of the RBD domains to engage in the binding to a second ACE2-Fc domain.

### Pseudotyped virus and SARS-CoV-2 virus neutralization assays show that dimeric ACE2-Fc can efficiently block viral entry

To translate these studies to cells, we compared the ability of ACE2 monomer and ACE2-Fc to neutralize SARS-CoV-2 pseudotyped virus to infect cells. Pseudotyped HIV-1 particles carry the wildtype SARS-CoV-2 Spike protein and are capable of delivering a NanoLuc luciferase reporter gene to ACE2-expressing HEK293T cells. Cells and pseudovirus were pre-incubated with serially diluted ACE2-Fc or monomeric ACE2, and luciferase activity was measured in cell lysates at 48 hours post infection. ACE2-Fc neutralized the SARS-CoV-2 pseudotyped particles at an IC_50_ = 0.96 μg/mL (4.75 nM), while ACE2 monomer did not substantially neutralize at the concentrations we tested, up to 48 μg/mL (237.5 nM) **(Figure 4A, 4B)**. This stark difference in efficacy between monomeric and dimeric ACE2 confirms that ACE2 monomer binds poorly to Spike.

**Figure 4.**
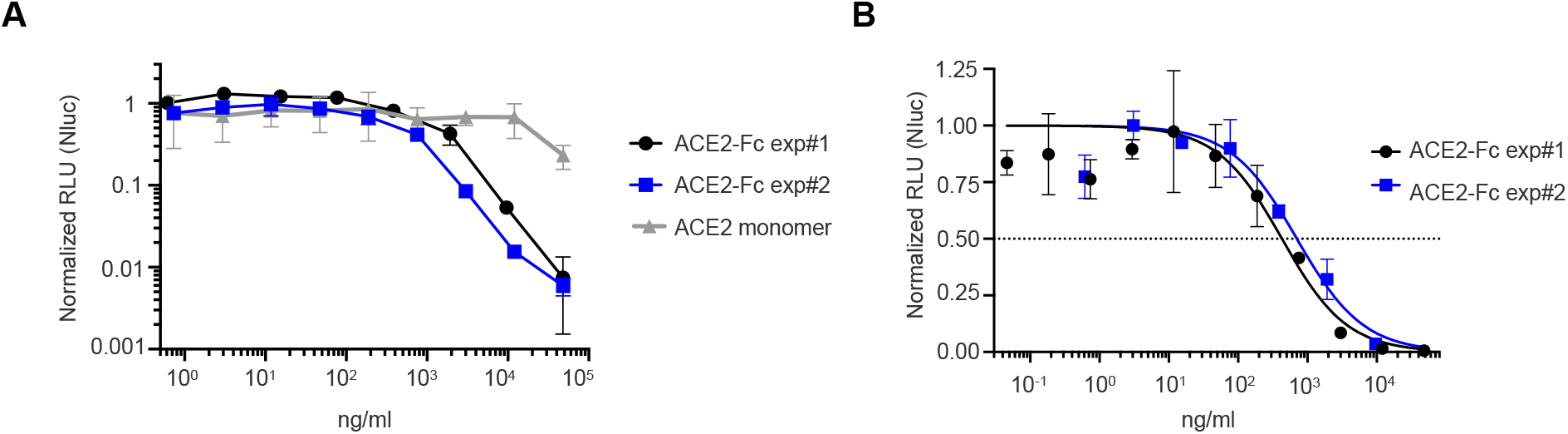
ACE2-Fc neutralizes pseudotyped virus more potently than ACE2 monomer. **(A)** Neutralization of luciferase-encoding pseudotyped lentivirus expressing SARS-CoV-2 Spike glycoprotein. Pseudotyped virus pre-incubated with ACE2-Fc or ACE2 monomer at indicated concentrations was used to infect HEK293T cells overexpressing ACE2. Luciferase activity in cell lysates were determined at 48 hours post infection and normalized to no treatment control. **(B)** The half-maximal inhibitory concentration (IC_50_) was obtained for the two ACE-Fc treatment by fitting the normalized data to a 4-parameter nonlinear regression. The average IC_50_ between two two biological replicates of ACE2-Fc treatment was 0.98 μg/ml (4.8 nM).

We further tested the ability of ACE2-Fc and ACE2 monomer to neutralize SARS-CoV-2 virus. SARS-CoV-2 live virus was pre-incubated with 100 nM of ACE2-Fc or ACE2 monomer prior to infecting VeroE6 cells, a monkey kidney epithelial cell line broadly used for studying viral infectivity. 16 hours post infection, cells were lysed and intercellular viral and host RNA was isolated, converted to cDNA, and quantified by qPCR. Consistent with the pseudotyped virus neutralization results, ACE2-Fc neutralized SARS-CoV-2 much more potently than ACE2 monomer **(Figure 5A, 5B**).

**Figure 5.**
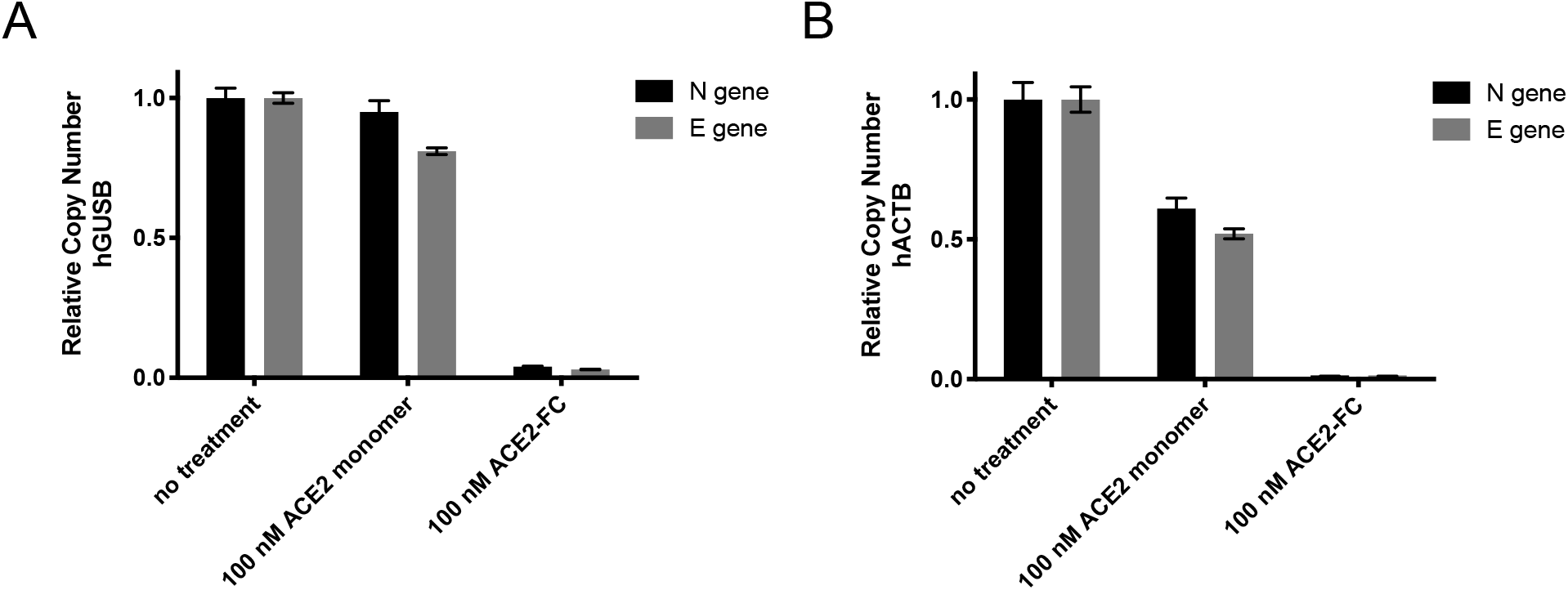
ACE2-Fc neutralizes SARS-CoV-2 live virus more potently than ACE2 monomer. SARS-CoV-2 live virus pre-incubated with 100 nM of ACE2-Fc or ACE2 monomer was used to infect VeroE6 cells. 16 hours post infection, cells were lysed and intercellular RNA was isolated and converted to cDNA. Viral entry and cellular transcription of viral genes was measured by qPCR. Relative copy number of viral N or E gene transcript for each treatment arm was determined using either **(A)** host gene GUSB (hGUSB) or **(B)** host gene ACTB (hACTB) as the reference control. Primers are cross reactive between *Homo sapiens* and *Cercopithecus aethiops* (VeroE6 cells).

## Discussion

Worldwide efforts are currently underway to develop effective and fast-acting clinical interventions to control the spread and mortality of SARS-CoV-2. While it is clear that the interaction between ACE2 and Spike-RBD plays a crucial role in viral entry, our current understanding is insufficient for the design of maximally effective therapeutic options. In this work, we systematically interrogated how an ACE2 dimer interacts with SARS-CoV-2 Spike trimer to understand the fundamental avidity properties of the Spike/ACE2 interaction.

The results from the BLI experiments, split reporter assays, and virus neutralization assays provide important insight into the conformational landscape of the RBDs in SARS-CoV-2 Spike and are summarized in **Figure 6A**. The decreased *k*_on_ of ACE2-Fc suggests the majority of RBDs in FL-Spike are in an ACE2-inaccessible, “down” conformation, which requires opening up for binding ACE2 (**Figure 6A**). The poor binding of monomeric ACE2 (**Figure 6B**) and strong binding to dimeric ACE2-Fc (**Figure 6C**) indicate intra-Spike avidity and ACE2 rebinding plays an important role in promoting this interaction. However, while we observed a highly productive avidity effect for the Spike-RBD-Fc/ACE2-Fc interaction, we did not observe the massive two-on-two avidity for the FL-Spike/ACE2-Fc interaction (**Figure 6D**). This weaker interaction could be due to RBD conformations in Spike, or a sub-optimal geometry of the two proteins for high-avidity binding. Nonetheless, we show that two or more of the RBD domains can be in the “up” conformation allowing binding of two ACE2-Fc on one Spike trimer (**Figure 6E**). This is evidenced in our BLI experiment where once bound to FL-Spike, the presence of the other RBDs slows down the dissociation of ACE2-Fc (**Figure 2F** in comparison to **2D**). This is further shown in the split-NanoLuc reporter experiments where ACE2 domains from two separate ACE2-Fc split-reporters can simultaneously bind a Spike trimer to generate an active luciferase (**Figure 3**). Together these results indicate that trimeric SARS-CoV-2 Spike interacts with ACE2 with limited but important intra-Spike avidity.

**Figure 6.**
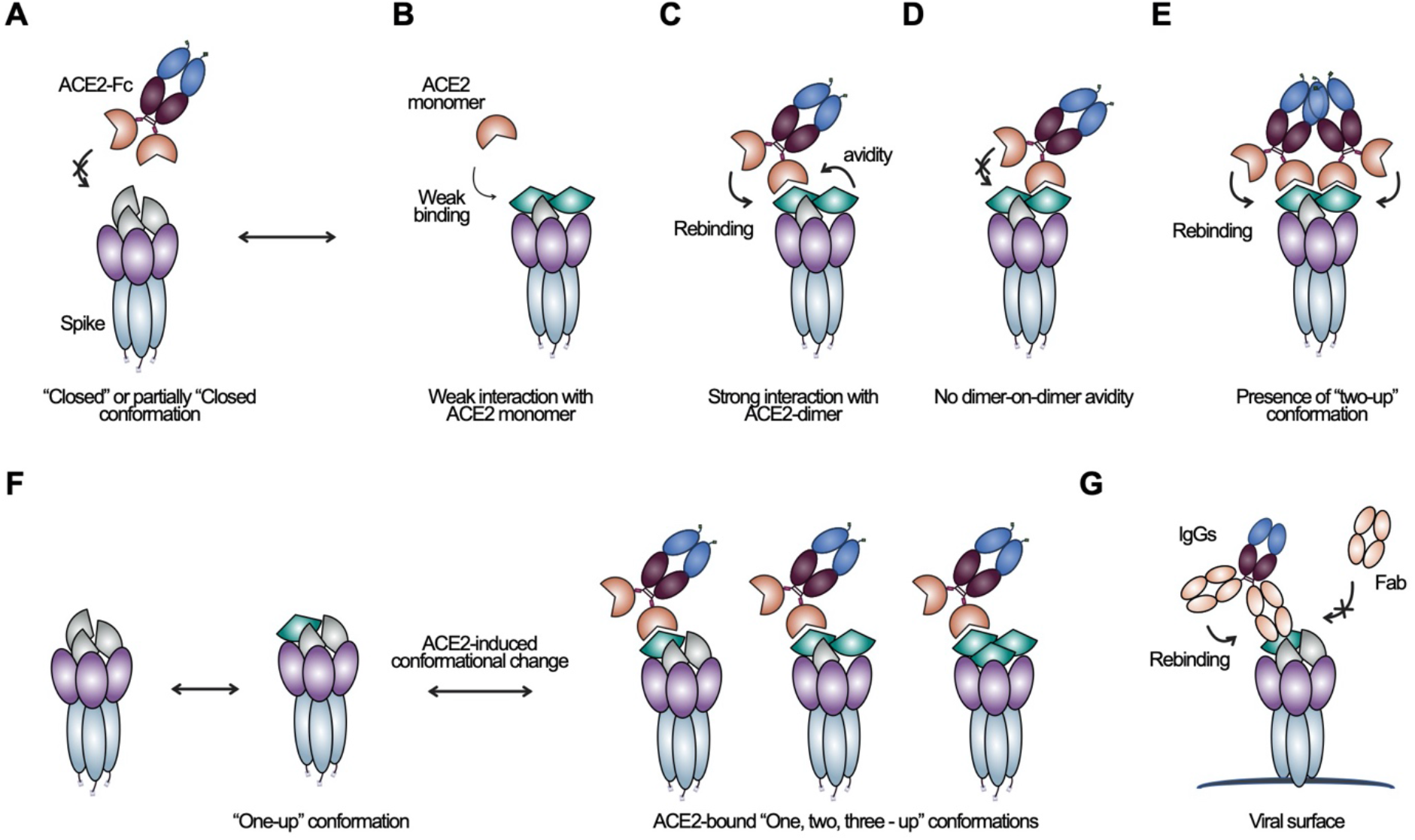
Model of the ACE2/Spike interaction and therapeutic strategies. **(A)** The majority of FL-Spike proteins are in the “closed” or partially “closed” conformation and “opening up” of RBD on Spike is necessary for ACE2 binding. **(B)** Monomeric ACE2 binds poorly to trimeric Spike, **(C)** ACE2-Fc binds much stronger to Spike than monomer, indicating that rebinding and intra-Spike avidity contribute significantly to the high-affinity nature of this interaction. **(D)** Dimeric ACE2-Fc does not interact with FL-Spike with a full two-on-two intramolecular avidity. **(E)** More than one RBD can be in the “up” conformation, enabling the engagement of separate ACE2 molecules. For **(B-E)**, only the “two-up” conformation is shown but other RBD conformations are also possible. **(F)** A proposed model where ACE2 binding may induce a conformational change in Spike, resulting in “two-up” or “three-up” RBD conformations. **(H)** Dimeric molecules such as ACE2-Fc or IgG will be more potent than a monomeric inhibitor for neutralizing SARS-CoV-2 virus. For all panels, RBD in the “open” conformation is colored in green and RBD in the “closed” formation colored in grey.

In addition to intra-Spike avidity, inter-Spike avidity could contribute to the improved affinity of ACE2-Fc especially in the intact virus. It is possible that ACE2-Fc could bridge two Spike molecules on the viral surface and bind with increased avidity, similar to what we observed in the high-density loading BLI experiment (**Supplementary Figure 3A**). Previous structural studies of SARS-CoV-1 shows the Spike proteins are densely distributed (Lin et al., 2004), with ~100 Spike molecules displayed on a viral particle with a diameter of ~100 nm (Beniac, Andonov, Grudeski, & Booth, 2006; Neuman et al., 2006). Using values from these studies, we estimate an inter-Spike distance of ~180 Å. The structure of the native ACE2 dimer (PDB: 6M17) shows that the two ACE2 arms (~80 Å apart) would not be able to span this predicted inter-Spike distance. An ACE2 monomer fused Fc with flexible linker (ACE2-Fc) may be sufficient to only bridge two Spikes that are <150 Å apart. Structural studies of IgGs have shown the antigen binding sites in the Fab arms on a flexible-hinge region of an Fc to be roughly ~117-134 Å apart (Sosnick et al. 1992). This would still fall short of the inter-Spike distance. However, these estimates are based on a rigid membrane where Spike proteins are uniformly distributed and also do not account for heterogeneity in Spike presentation across viral particles. This may not be the case as coronaviruses are an enveloped virus with a fluid lipid membrane that allows for Spike protein mobility and clustering, and potentially enable inter-Spike binding by ACE2 dimers or IgGs. Indeed, cryo-EM images of SARS-CoV-1 show irregularity and lack of symmetry of Spike distribution on the coronavirus envelope (Beniac et al., 2006). As there are no direct reports of inter-Spike avidity for dimeric ACE2, further experiments are needed to thoroughly examine the Spike/ACE2 interaction on the viral surface.

While all cryo-EM structural studies to date on SARS-CoV-2 Spike have identified only the “closed” or “one-up” RBD conformation (Cao et al., 2020; Pinto et al., 2020; Walls et al., 2020; Wrapp, Wang, et al., 2020), our split reporter assay identified a population of Spike in “two-up” or “three-up” conformation. The relative population of these RBD conformers and whether they exist in an ACE2-unbound state or emerge only upon ACE2 binding remains unknown (**Figure 6F**). Previous studies have suggested that ACE2 binding could lead to conformational change in SARS-CoV-1 or 2 Spike (Song et al., 2018; Yuan et al., 2020). Molecular dynamic simulation of SARS-CoV-2 Spike bound to ACE2 found that there is significant flexibility in the RBD conformation (Peng et al., 2020), and an EM study of SARS-CoV-1 Spike reported that the distribution of the RBD conformers was very different in the ACE2-bound structure compared to the unbound structure (Kirchdoerfer et al., 2018). In accordance with these studies, our results support the model that ACE2 binding induces a conformational change in Spike and enables two or more RBDs to be in the “up” conformation (**Figure 6F**). This complex and dynamic nature of the Spike/ACE2 interaction is likely to play a key role in the biology of SARS-CoV-2 infection. We anticipate that further characterization of the interaction between SARS-CoV-2 Spike and native ACE2 will elucidate the exact nature of this binding.

Furthermore, our findings have important ramifications in developing a successful protein therapeutic for COVID-19. Recombinant ACE2 proteins or antibodies which can block the host ACE2-viral Spike interaction are promising as anti-SARS-CoV-2 therapeutics. The much higher potency of ACE2-Fc in comparison to ACE2 monomer we observed in the live virus neutralization assays suggest that monovalent therapeutics such as ACE2 monomers or Fab domains will likely be much less effective than multimeric formats such as dimeric ACE2 ectodomain (Monteil et al., 2020), ACE2-Fc (Lei et al., 2020; Yan et al., 2020) or IgG (**Figure 6G**). Finally, our results highlight the importance of understanding the ACE2-Spike interaction in the context of the FL-Spike trimer. We also hope that the Spike and ACE2 constructs generated here are useful tools to the greater research community to enable better understanding of SARS-CoV-2 biology.

## Acknowledgements

We thank members of the Wells Lab, particularly those working on the Covid-19 project for their efforts and contributions. We also thank Dr. John Pak (Chan Zuckerberg Initiative Biohub) and Dr. Florian Krammer (Mt. Sinai Icahn School of Medicine) for providing plasmids, and Dr. Abigail Elizabeth Powell from Dr. Peter Kim’s laboratory (Stanford University) for advice on FL-Spike expression. We also acknowledge Dr. Pamela Bjorkman (Caltech University), and Dr. Joel Ernst and Dr. Raul Andino (UCSF) for insightful discussions and feedback.

## Author Contributions

I.L., S.K.E., S.A.L. performed all experiments except those explicitly stated. X.X.Z, L.L.K., J.A.G. designed and cloned the ACE2-Fc-reporter fusions. X.X.Z, I.L., S.A.L., J.L, K.K.L. and J.A.W. designed the research and analyzed the data for the Biolayer interferometry experiments. S.K.E. X.X.Z, and J.A.W. designed the research and analyzed the data for the split reporter experiments. B.S.Z, P.S., N.J.R, S.A.L., O.S.R. designed and/or conducted the live virus neutralization assays. F.M., J.C.L., F.S., Y.W., D.F.R., M.C.N, T.H., P.B., designed and/or conducted the pseudotyped virus neutralization assays. X.X.Z, S.A.L., L.L.K., S.K.E., I.L., K.K.L and J.A.W. co-wrote the manuscript.

## Materials and Methods

### Plasmids construction

Plasmids were constructed by standard molecular biology methods. The FL-Spike plasmid was a generous gift from the Pak lab (Chan Zuckerberg Initiative Biohub) and Krammer lab (Icahn School of Medicine at Mount Sinai). The DNA fragments of Spike-RBD, ACE2 and LgBiT were synthesized by IDT Technologies. The Spike-RBD-TEV-Fc-AviTag, ACE2-TEV-Fc-AviTag, Spike-RBD-8xHis-AviTag, ACE2-8xHis-AviTag plasmids were generated by subcloning the Spike-RBD or ACE2 DNA fragment into a pFUSE-hIgG1-Fc-AviTag vector (adapted from the pFUSE-hIgG1-Fc vector from InvivoGen). The ACE2-Fc-LgBiT fusion plasmids were generated by subcloning the gene fragments of LgBiT to the N- or C-terminus of the ACE2-TEV-Fc-AviTag vector with a 10-amino acid (N-terminal fusion) or 5-amino acid (C-terminal fusion) linker. The SmBiT tag in the ACE2-Fc-SmBiT fusion plasmids was generated by overlap-extension PCR, which also has a 10-amino acid (N-terminal fusion) or 5-amino acid (C-terminal fusion) linker to the ACE2 or Fc domains. The C-terminal AviTag was removed from all the ACE2-Fc reporter plasmids. Complete plasmid sequences are available upon request.

### Expression and purification of ACE2 and Spike constructs

The ACE2 and Spike proteins were expressed and purified from Expi293 BirA cells according to established protocol from the manufacturer (Thermo Fisher Scientific). Briefly, 30 μg of pFUSE (InvivoGen) vector encoding the protein of interest was transiently transfected into 75 million Expi293 BirA cells using the Expifectamine kit (Thermo Fischer Scientific). Enhancer was added 20 h after transfection. Cells were incubated for a total of 3 d at 37 °C in an 8% CO2 environment before the supernatants were harvested by centrifugation. Fc-fusion proteins were purified by Protein A affinity chromatography and His-tagged proteins were purified by Ni-NTA affinity chromatography. Purity and integrity were assessed by SDS/PAGE. Purified protein was buffer exchanged into PBS and stored at −80 °C in aliquots.

### Generation of ACE2 monomer

ACE2 monomer was obtained by TEV treatment of ACE2-Fc and subsequent purification. 50 μl Ni-NTA agarose (Qiagen) and 50 μl Neutravidin resin (Thermo Fisher Scientific) were washed with PBS-25 mM imidazole twice and combined in 100 μl PBS-25 mM imidazole. Next, 20 μg His-Tagged recombinant TEV protease and 1 mg purified ACE2-Fc protein were mixed, and the reaction tube was rotated at 4 °C for 30 minutes. The cleavage reaction was then incubated with the washed beads, rotating, at 4 °C for 30 minutes. While the incubation occurred, an additional 25 μl of magnetic Protein A beads and 25 μl or Ni-NTA beads were prepared as described before. Supernatant from the first bead clearance was transferred to the newly prepared beads and allowed to incubate for an additional 30 minutes at 4 °C. To remove beads from the protein supernatant, reaction mixture was spin filtered at 1000 g for 2 min and washed with an additional 250 uL of PBS-25 mM imidazole. The His-tagged TEV, biotinylated Fc, and the uncut ACE2-Fc remained on the beads while the monomeric ACE2 was isolated in the flow-through. The purity of monomeric ACE2 was confirmed by SDS-PAGE electrophoresis. Purified protein was buffer exchanged to PBS and store at −80 °C in aliquots.

### Differential scanning fluorimetry

To assess the stability of proteins, we measured the melting temperature (T_m_) by doing differential scanning fluorimetry (DSF) as the method described previously (Hornsby et al., 2015). Briefly, purified protein was diluted to 0.5 μM or 0.25 μM in DSF buffer containing Sypro Orange 4x (Invitrogen) and PBS. 10 μL of reaction mixture was transferred to one well of a 384-well PCR plate. Duplicate was prepared as needed. In a Roche LC480 LightCycler, the reaction was heated from 30°C to 95°C with a ramp rate of 0.3°C per 30 sec. The intensities of the fluorescent signal at an ~490 nm and ~575 nm (excitation and emission wavelengths) were continuously collected. The curve peak corresponds to the melting temperature of the protein. Data was processed and T_m_ was calculated using the Roche LC480 LightCycler software.

### *In vitro* binding experiments

Biolayer interferometry data were measured using an Octet RED384 (ForteBio). Biotinylated Spike or Spike RBD protein were immobilized on the streptavidin (SA) biosensor (ForteBio). After blocking with biotin, purified ACE2 proteins in solution was used as the analyte. PBS with 0.05% Tween-20 and 0.2% BSA was used for all diluents and buffers. A 1:1 monovalent binding model was used to fit the kinetic parameters (*k*_on_ and *k*_off_).

### Magnetic bead and solution based NanoBiT assays

For the Spike-Fc magnetic bead assay, magnetic beads were prepared by taking 100 μL of Streptavidin Magnesphere Paramagnetic Particles (Promega) and incubated with 5 μM of Spike-Fc-AviTag for 30 minutes, rotating at room temperature. Following, the beads were blocked with 10 uM biotin for 10 minutes. The beads were washed three times with PBS + 0.05% Tween + 0.2% BSA. 10 μL of 10-fold dilutions of the beads were incubated with 10μL of premixed 2 nM ACE2-Fc-SmBiT and ACE2-Fc-LgBiT fusions. The sample was incubated shaking at room temperature for 20 minutes. NanoGlo Lucfierase substrate (Promega) diluted in NanoGlo Luciferase buffer was added to each well (15 μL) and luminescence was measured on a Tecan M1000 plate reader after 10 minutes. For the FL-Spike detection, 10 μL of FL-Spike dilutions were combined with 10 μL of premixed 2 nM SmBiT-ACE2-Fc and LgBiT-ACE2-Fc. Samples were incubated with substrate and luminescence was detected as described above.

### Pseudotyped virus neutralization assay

HEK293T cells overexpressing full-length ACE2 carrying two inactivating mutations in the catalytic domain (H374N & H378N) were generated using standard lentivirus transduction. The cells were maintained in DMEM (Gibco) with 10% heat-inactivated fetal bovine serum, Gentamycin and 5 μg/ml Blasticidin.

Pseudotyped HIV-1 particles expressing the SARS-CoV-2 Spike glycoprotein and NanoLuc luciferase as a reporter were generated by transfection of HEK293T cells with pNL4-3DEnv-NanoLuc and pSARS-CoV2-S_trunc_. pNL4-3DEnv-NanoLuc was derived from pNL4-3 (Adachi et al., 1986) by inserting a 940 bp deletion after the *vpu* stop-codon, resulting in loss of Env-expression. The NanoLuc Luciferase reporter gene (*Nluc*, Promega) was inserted in place of bp 1-100 in the *nef*-frame. pSARS-CoV2-S_trunc_ was generated by insertion of a human-codon optimized gene encoding for 19 AAs C-terminally truncated SARS-CoV-2 Spike (Geneart) into pCR3.1.

Supernatants containing virus were harvested and filtered 48 hr post transfection and used for infection of ACE2-overexpressing 293T cells. Pseudotyped virus was pre-incubated with serially diluted ACE2-Fc or ACE2 monomer at 37 °C for 1 hr before addition to cells., Cells were washed twice with PBS 48 hr post infection and lysed with Luciferase Cell Culture Lysis 5x reagent (Promega). Relative luminescence units were normalized to those derived from cells infected with SARS-CoV-2 pseudotyped virus in the absence of Ace2-Fc/Ace2 monomer. The half maximal inhibitory concentration (IC_50_) was determined using 4-parameter nonlinear regression (GraphPad Prism). Each concentration of Ace2-Fc/Ace2 monomer was tested in duplicate and reported as an average and standard deviation for each experiment. Two biological replicates of ACE2-Fc treatment were obtained (exp #1, exp #2).

### SARS-CoV-2 virus neutralization assay

All handling and experiments using SARS-CoV-2 was conducted under Biosafety Level 3 containment with approved BUA and protocols. SARS-CoV-2 clinical isolate 2019-nCoV/USA-WA1/2020 was obtained through BEI Resources (Harcourt et al., 2020). Prior to experiments, virus was passed in Vero E6 cells to create working stocks and titers were measured by plaque formation assay. For experiments, Vero E6 cells were cultured in Minimal Essential Media (MEM), 10% Fetal Bovine Serum (FBS), 1% Pen-Strep and seeded on 6-well culture plates at 3.8E5 cells/well the day prior. Infection with SARS-CoV-2 was performed using MOI of 0.1. Virus was incubated in infection media (EMEM 0% FBS) containing 100 nM monomeric ACE2, 100 nM dimeric ACE2-Fc, or no blocker for 1 hour at 37 °C. Culture media was removed from Vero E6 cells and 300 μL of the blocker/virus inoculum was added for 1 hour at 37 °C. Subsequently, 1 mL of cell culture media was added and cells were incubated at 37 °C for an additional 16 hours before RNA harvest.

Viral entry into cells and cellular transcription of viral genes was measured by qPCR. Cellular RNA was isolated and converted to cDNA using RNAeasy RNA extraction kit (Qiagen) and Quantitect Reverse-transcriptase kit (Qiagen) according to manufacturer instructions. qPCR reactions were prepared using SYBR Select Master Mix (Thermo) and the following conditions: for N gene, E gene, and hGUSB gene primer (cross reactive with *Cercopithecus aethiops*) concentration was 400 nM and annealing temperature was 58 °C, and for hACTB gene primer (cross reactive with *Cercopithecus aethiops*) concentration was 500 nM and annealing temperature was 60 °C. Primer sequences (IDT) were the following – viral genes: N_F = CACATTGGCACCCGCAATC; N_R = GAGGAACGAGAAGAGGCTTG; E_F = ACAGGTACGTTAATAGTTAATAGCGT; E_R = ATATTGCAGCAGTACGCACACA; and host genes: hGUSB_F = CTCATCTGGAATTTTGCCGATT; hGUSB_R = CCGAGTGAAGACCCCCTTTTTA; hACTB primers were IDT PrimeTime assay reagent. Relative copy number of viral transcript level compared to cellular transcript was determined using the ΔΔCT method.

---

**Supplementary Figure 1.**
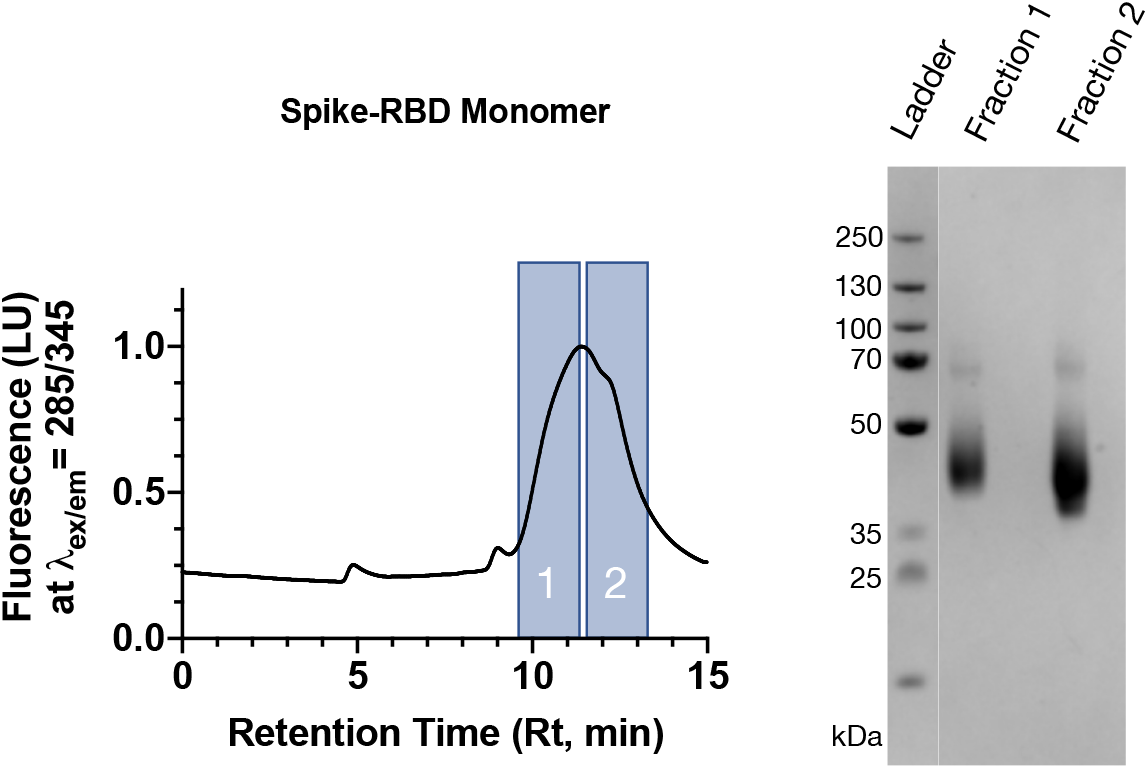
SDS-PAGE analysis of SEC fractions shows pure Spike-RBD protein at the correct molecular weight.

**Supplementary Figure 2.**
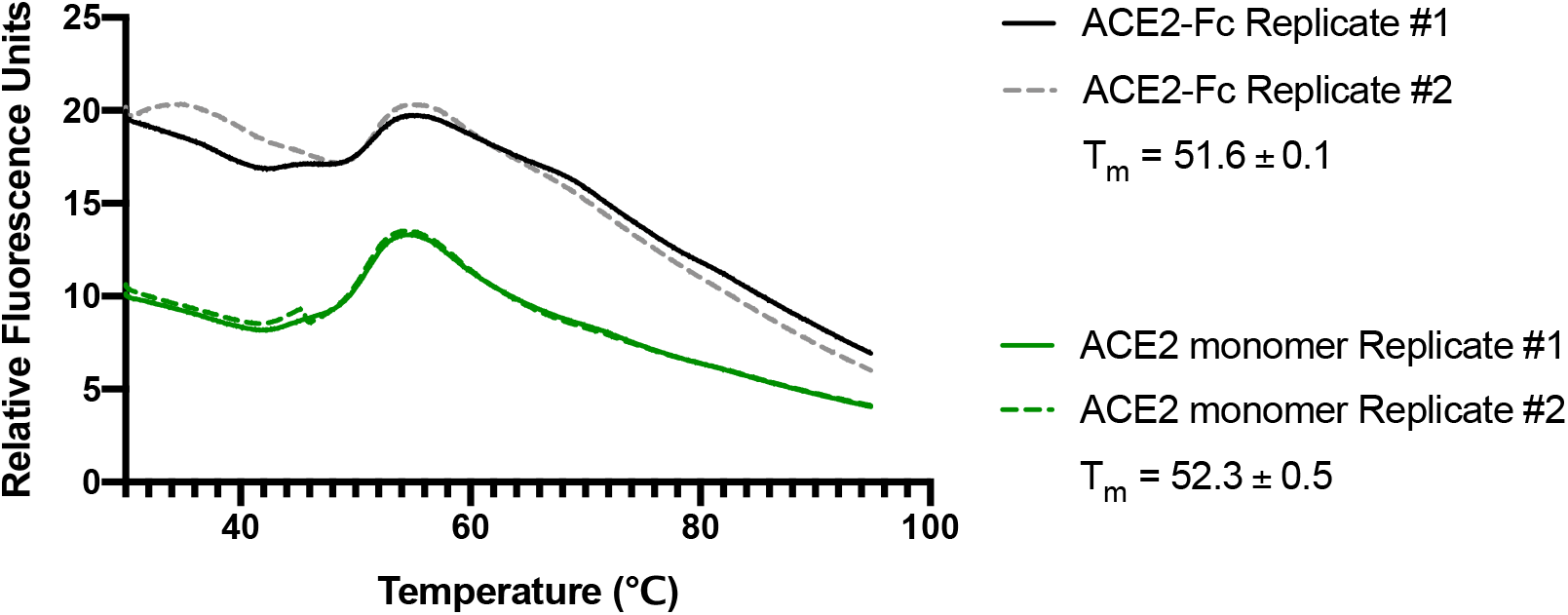
Melting temperature of reagents measured by Differential Scanning Fluorimetry (DSF). T_m_ is reported as an average of two replicates.

**Supplementary Figure 3:**
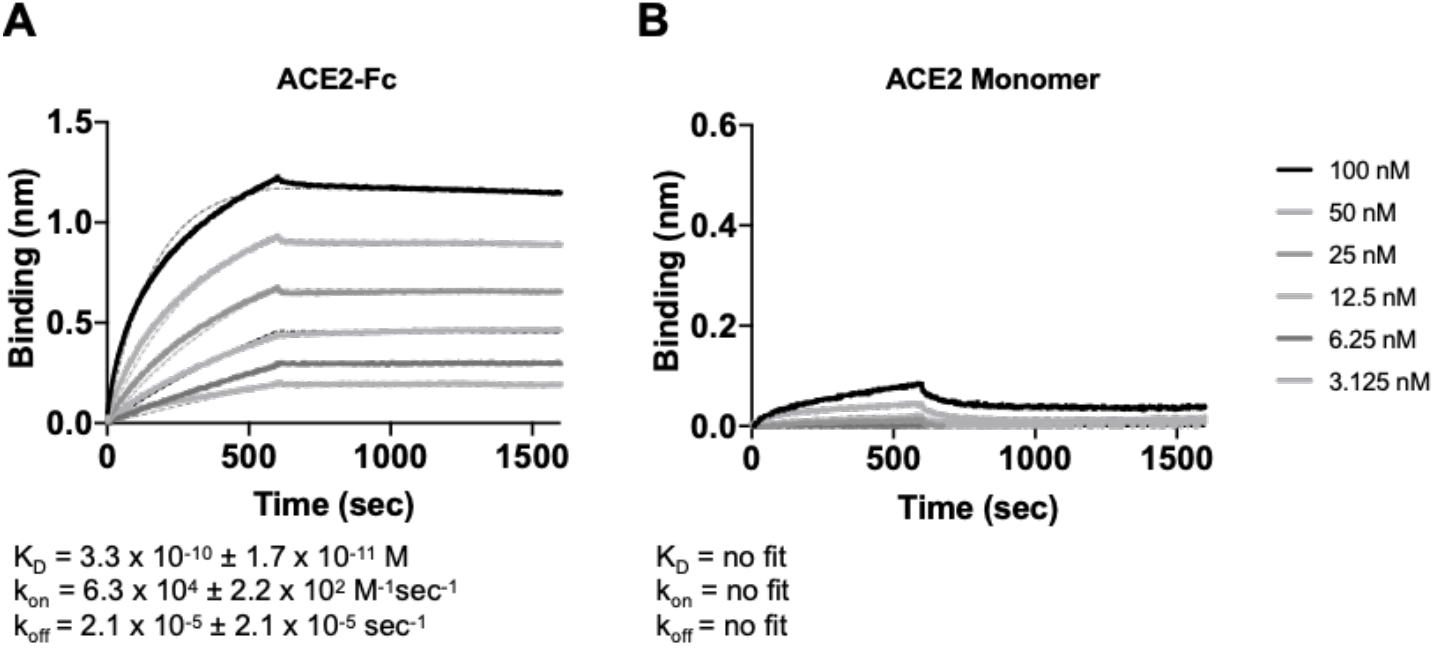
Saturating FL-spike on biosensor reveals intermolecular avidity of ACE2-Fc. FL-Spike was loaded onto sensors until saturation. **(A)** ACE2-Fc was used as analytes in solution, with concentrations beginning at 100 nM to 3.125 nM at 2-fold dilutions. The affinities for the ACE2 dimeric to saturating trimeric FL-Spike was 0.33 nM, or 18-fold higher in affinity compared to non-saturating conditions (5.9 nM, **Figure 2F**). **(B)** ACE2 monomer did not bind FL-Spike strongly even when FL-Spike was loaded until saturation.

**Supplementary Figure 4.**
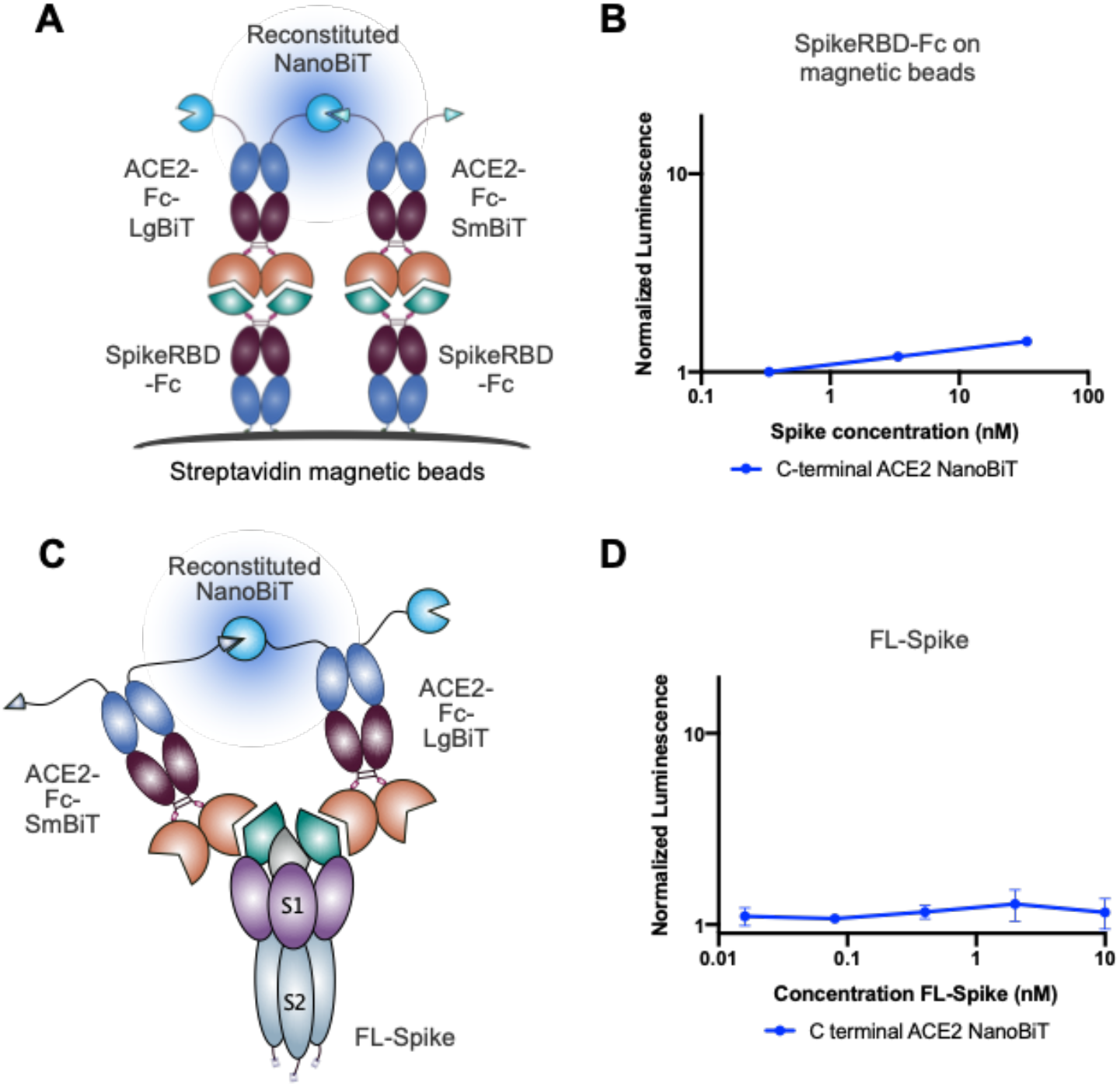
C-terminally fused ACE2-Fc Split Luciferase Reporter is not as sensitive as the N-terminal fusions. **(A)** Cartoon depiction of the C-terminal ACE2 NanoBiT sensor system to detect Spike-RBD-Fc bound to streptavidin magnetic beads. **(B)** The C-terminal ACE2 NanoBiT system is able to detect Spike-RBD-Fc bound magnetic beads, but with lower sensitivity than the N-terminal fusions. **(C)** Cartoon depiction of the C-terminal ACE2 NanoBiT sensor system to detect FL-Spike in solution. **(D)** The C-terminal ACE2 NanoBiT system is not able to detect Spike-RBD-Fc bound magnetic beads due to lower sensitivity. For all assays, the luminescence signal is normalized to a no-bead control and the average and standard deviation is plotted (N=3).

